# Effects of TeaCrine® (theacrine), Dynamine™ (methylliberine), and caffeine on gamer psychomotor performance in a first-person shooter video game scenario

**DOI:** 10.1101/2021.04.21.440794

**Authors:** Michael B. La Monica, Jennifer B. Listman, Ian Donovan, Taylor E. Johnson, David J. Heeger, Wayne E Mackey

**Author notes:** Corresponding author: Michael B. La Monica.

## Abstract

**Purpose:** To compare the effects of purported cognitive enhancing dietary bioactive ingredients on subjective and objective measures of cognitive and motor performance during a first-person shooter video game.

**Methods:** Using a placebo-controlled crossover design, nine healthy men (23.4±5.7 yr, 178.9±5.8cm, 86.0±17.1kg) completed four 20-minute gaming sessions designed to assess cognitive, motor, and perceptual skills via artificial intelligence-driven battery of tasks (Aim Lab). Participants ingested either a placebo (PL), caffeine (CAFF), or caffeine + methylliberine (Dynamine™) + theacrine (TeaCrine®) (CMT). Before and after each gaming session participants rated various feelings of affect. Data were analyzed using mixed factorial ANOVA, bootstrapping post-hoc tests with 95% confidence intervals, and effect sizes.

**Results:** Compared to PL, self-assessed performance was significantly increased in CMT vs. PL (p=0.035) and self-assessed jitteriness was increased by CAFF vs. PL (p=0.043). CMT was associated with a greater improvement of participants’ visuo-spatial working memory from baseline vs. PL (p=0.04) and CAFF (p=0.033). CAFF had a greater decrease in reaction time for false alarms (indicating diminished cognitive control) from baseline vs. PL (p=0.002) and CMT (p=0.001) and a greater increase for time on target tracking vs. PL (p=0.008) and CMT (p=0.047). Compared to PL, CMT was associated with a greater decrease in median kill time (indicating improved speed) (p=0.017). Compared to PL, systolic blood pressure was significantly increased by CAFF (p=0.025) and CMT (p=0.020) but remained within normal limits.

**Conclusions:** Acute CMT supplementation improved cognitive and motor abilities in recreational gamers. The addition of theacrine and methylliberine to caffeine may lessen some undesirable effects of isolated caffeine ingestion on cognitive control and jitteriness.

## Introduction

Video games in the US are a $67-billion-dollar industry with an average growth of 14.7% over the past five years (2015-2020)^1^. The worldwide esports market brought in just over 1 billion dollars in 2020^2^. The popularity among individuals playing games has been on the rise with 64% of adults over the age of 18 playing video games and 75% of households having at least one video game player currently^3^.

Video games encompass a variety of cognitive tasks that challenge players in all dimensions of their intellectual capabilities. More specifically, video games involve attention, visuospatial tasks, cognitive workload, specialized psychomotor behavior skill acquisition, and cognitive control^4^. In other words, successful players must have attention control, the ability to recognize, perceive, and manipulate visual stimuli, visuomotor coordination and navigational skills^4^. Players must also demonstrate cognitive control abilities such as reactive and proactive inhibition, task switching, and working memory^5^. High level gamers have characterized successful performance as having the ability to think strategically, while making quick smart decisions, not dwelling on past events, avoiding distractions, maintaining focus, and retaining a positive attitude^6^. Video games themselves can boost visual attention^7^. In fact, action video games have been shown to enhance an individual’s ability to track multiple objects^7,8^ while first person shooters may enhance reaction times and processing speed^9^.

Cognitive enhancing dietary supplements (i.e., nootropics) have been used by esports athletes to gain a competitive advantage and given their increasing popularity among esports athletes, there is a clear need to vet products for safety and efficacy^10^. Caffeine, a methylxanthine, has shown to improve mental performance (i.e., faster reaction times and motor performance)^11^. Caffeine can be beneficial in enhancing reaction time, vigilance, and attention and may mitigate low levels of awareness and activation during prolonged monotonous activities^12^. Yuan et al.^13^ found that coffee enhanced response time and accuracy in the Stroop test. Caffeine has also been shown to enhance the ability to inhibit inappropriate responses (i.e., false alarms when being faced with incompatible stimuli) in habitual users^14^, a function important in video game performance, which often incorporates reactive inhibition. Rodgers et al.^11^ showed 150 mg of caffeine enhanced motor speed and simple reaction time during simple computer tasks but did not enhance recognition memory showing that caffeine made them faster not smarter. However, not all studies have found benefits. Thomas et al.^15^ demonstrated that a multi-ingredient energy drink containing 150 mg of caffeine did not enhance cognitive performance or motor speed before or after a competitive gaming series. Nonetheless, because caffeine offers a potential advantage to gamers its use is enticing.

Compounds with structural similarities to caffeine, such as methylurate purine alkaloids may provide additional benefits in comparison to caffeine alone. One of these compounds is TeaCrine® (theacrine), a methylurate which has been shown to promote increased levels of energy and reduced fatigue^16^. The half-life of caffeine (6.2 ± 3.8hr) and TeaCrine® (26.1 ± 13.7hr) are different, but the co-administration of the two altered TeaCrine® pharmacokinetics enhancing the exposure and area under the curve^17^. TeaCrine® appears to provide a mild stimulant and calming effect acting through the adenosine and dopamine systems^18^, thereby potentially allowing for a smaller dose of caffeine which can negate the potential side effects in caffeine sensitive individuals. Ziegenfuss et al.^19^ showed increased energy, focus, lower anxiety, increased motivation to exercise and concentration without jitteriness during acute supplementation with TeaCrine®. TeaCrine® has been shown to be safe after eight weeks of use^20^ and does not appear to exhibit a tachyphylaxis or habituation effect like caffeine, likely due to different structure-activity relationships and allosteric modulation at the adenosine and dopamine receptors^18^. As opposed to either supplement alone, TeaCrine® and caffeine may work better together to enhance energy, mood, and focus^21^. Synergistically, caffeine and TeaCrine® have been shown to lower jitteriness compared to caffeine alone, while the two compounds together did not elevate blood pressure or heart rate^21^.

Another compound hypothesized to have similar physiological and cognitive effects as caffeine and TeaCrine® is methylliberine, also known as Dynamine™. Specifically, methylliberine is a methoxyurate, and like theacrine is also found in plants of the genus *Coffea, Theobroma* and *Ilex* as a downstream metabolite of caffeine via a theacrine intermediate^22^. Minimal human research has been done with methylliberine other than demonstrating the safety of acute and long-term consumption in young healthy men and women^23,24^ and modeling its cellular absorption^25^. Methylliberine and caffeine may have a synergistic relationship to enhance mood^23^. Together, all three compounds (i.e., TeaCrine®, Dynamine™, and caffeine) have been shown to be safe together or independently^23,24^. Since caffeine may augment the effects of TeaCrine®, co-administration of all these substances may further enhance cognitive activity and control in a computer gaming task. Finally, because these compounds have different cellular absorption rates^25^ and half-lives^26^, performance benefits may be maintained for a longer duration than caffeine itself. However, these hypotheses remain untested.

The primary aims of this study were to measure and compare changes in psychometric indices of energy, alertness, focus, creativity, decision making ability, processing speed, irritability, and jitteriness, and to objectively assess and compare speed, precision, accuracy, reaction time, visuo-spatial working memory capacity, and cognitive control during a first person-shooter gaming scenario, and to assess safety and tolerability as determined by vital signs and adverse events. Specifically, we compared caffeine alone and caffeine plus TeaCrine® and Dynamine™ to a placebo.

## Methods

### Experimental Design

This was a repeated measures crossover design in which participants visited the lab on seven occasions (one screening visit, three familiarization visits, and three testing visits). During the initial screening visit each participant’s medical history and blood work (CBC, CMP, and lipid panel) were assessed, and baseline diet was evaluated. During the next three visits (i.e., familiarization trials) participants had their vitals measured before a gaming session (i.e., a playlist of tasks specially designed to measure and assess cognitive, motor, and perceptual skills) and filled out visual analog scales (VAS) before and after a single 20-minute gaming session. During the final three visits (i.e., testing trials) subjects performed the same measures as the familiarization trials but repeated them at specific intervals with respect to the treatment conditions [i.e., 0 (pre-treatment), 60 min, 120 min, and 180 min post treatment]. During the testing visits each participant ingested one of three test products after their baseline (0 min) gaming session. All study procedures were conducted in accordance with the Declaration of Helsinki and the International Conference on Harmonization Guidelines for Good Clinical Practice (ICH-GCP). IRB approval for this study was approved by IntegReview, LLC (Austin, TX) on June 12, 2020.

### Participants

Nine healthy men completed all study visits. Each participant was a self-proclaimed recreational gamer. All participants were in good health as determined by physical examination and medical history, between the ages of 18 and 45 years, and had a body mass index (BMI) of 18.5-35 kg•m^-2^. Prior to participation, all participants indicated their willingness to comply with all aspects of the experimental and supplement protocol. Participants were excluded if they: (a) had a history of diabetes or pre-diabetes; (b) had a history of malignancy in the previous 5 years except for non-melanoma skin cancer (basal cell cancer or squamous cell cancer of the skin); (c) had prior gastrointestinal bypass surgery; (d) known gastrointestinal or metabolic diseases that might impact nutrient absorption or metabolism (e.g. short bowel syndrome, diarrheal illnesses, history of colon resection, gastro paresis, Inborn-Errors-of-Metabolism); (e) had any chronic inflammatory condition or disease; (f) had a known allergy to any of the ingredients in the supplement or the placebo; (g) had currently been participating in another research study with an investigational product or have been in another research study in the past 30 days; (h) had a caffeine intake of three or more cups of coffee or equivalent (>400 mg) per day; (i) were consuming energy drinks or pre-workout products, and taking stimulants or nootropic supplements 24 hours prior to enrollment and any testing visit to the lab; (j) used corticosteroids or testosterone replacement therapy (ingestion, injection, or transdermal); (k) had any other diseases or conditions that, in the opinion of the medical staff, could confound the primary endpoint or place the participant at increased risk of harm if they were to participate; or (l) did not demonstrate a verbal understanding of the informed consent document.

Participants were instructed to follow their normal diet and activity patterns during the study. Participants were required to complete a 24-hour diet record prior to arriving at the laboratory for their first initial screening visit. Participants were given a copy of this dietary record and instructed to duplicate all food and fluid intake 24 hours prior to their next laboratory visit. In addition to replicating food and fluid intake for 24 hours prior to their subsequent testing session, study participants were also asked to refrain from exercise 24 hours prior, abstain from caffeine 12 hours prior, and arrive 8 hours fasted to all testing sessions.

### Familiarization visits

Each participant began their visit by filling out a visual-analog scale (VAS) for subjective feelings of energy, alertness, focus, creativity, decision making ability, processing speed, irritability and jitteriness as well as measuring their vitals (i.e., heart rate and blood pressure). Next each participant completed a 20-minute gaming session on specifically designed gaming software (Aim Lab, Statespace) with a gaming computer, mouse, and headphones (ASUS). After the gaming session participants filled out another VAS rating their overall perceived gaming performance.

### Testing visits

Participants completed four intervals of the same procedures as the familiarization trials (i.e., VAS, vitals, 20-minute gaming session, VAS). Immediately after their first assessment participants ingested one of three different supplements. On these days additional 20-minute gaming sessions occurred at the end of hour 1 (40-60 min post-ingestion), hour 2 (100-120 min post-ingestion), and hour 3 (160-180 min post-ingestion)] (Figure 1).

**Figure 1.**
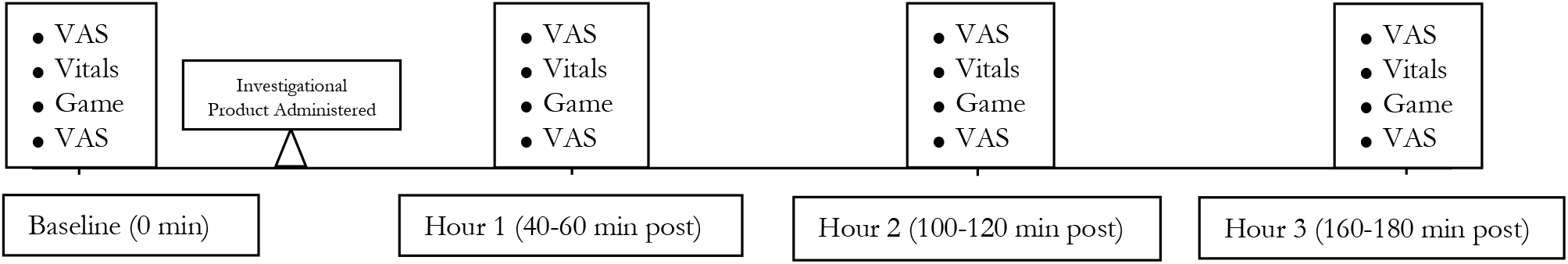
Study Timeline of Testing Visits

### Supplement Protocol

Throughout the study protocol, all supplements were prepared in capsule form for oral ingestion and packaged in coded generic containers for administration. Participants orally ingested 1) placebo (rice flour), 2) caffeine (125 mg), or 3) caffeine (125 mg) + 75 mg methylliberine (as Dynamine™) + 50 mg theacrine (as TeaCrine®). Each treatment was administered after the first gaming session during the final three visits (i.e., testing visits) in a double-blinded fashion using a Latin Square approach to minimize order effects. All trials were conducted at the research center under the supervision of a study team member. In addition, all supplements were tested by an independent third-party manufacturer for purity and potency.

### Anthropometric and Other Resting Measures

Standing height was determined using a wall-mounted stadiometer and body weight was measured using a Seca 767TM Medical Scale. Resting heart rate and blood pressure were measured using an automated blood pressure cuff (Omron HEM-780) before, 40, 100, and 160 minutes after ingestion of each assigned supplement.

### Visual-Analog Scales

100-mm anchored VAS were completed before, 40, 100, and 160 minutes after ingestion of each acute supplement. VAS were anchored with “Worst Possible” and “Lowest Possible” or “Best Possible” and “Highest Possible” and assessed subjective ratings of energy, alertness, focus, creativity, ability to make making decisions, level of processing speed, irritability, and jitteriness. An additional VAS that assessed their overall assessment of their gaming performance was administered immediately after each gaming session at the above stated intervals.

### Gaming software

Statespace (https://statespace.gg New York, NY) developed a first-person shooter (FPS) game, Aim Lab, based on sensorimotor neuroscience, psychophysics, and computational/theoretical neuroscience to assess and train players. Aim Lab is free and available to download on Steam (https://store.steampowered.com/app/714010/Aim_Lab/). Aim Lab was written in the C# programming language using the Unity game engine. Unity is a cross-platform video game engine used for developing digital games for computers, mobile devices, and gaming consoles. Aim Lab developed a “playlist” of video game tasks in a virtual environment that overlay a cognitive assessment.

The non-invasive, objective gamified assessment covers aspects of cognition, perception, and motor control addressed in traditional neuropsychological batteries such as working memory, cognitive control, processing speed, visual motion perception, ballistic movement control, tracking movement control, and divided attention. A short session of gameplay results in baseline objective, granular neuropsychological measurements for each individual. Repeated gameplay serves to monitor any changes in cognitive and motor performance.

Participants controlled their virtual weapon in the Aim Lab gamified assessment using a standard gaming mouse on a gaming computer. The assessment tool consisted of repeated rounds of four tasks, each of which is a skill assessment within the context of a FPS video game. Each task was tailored to a facet of FPS play, lasted between 1-2 minutes and simulated scenarios familiar to video game players. In total, the assessment took approximately 20 minutes per session, including brief breaks and reminders of instructions. The tasks and their order were predetermined and identical for all participants, across trials.

Data from each round of play in Aim Lab was stored in private cloud servers via Amazon Web Services (Seattle, WA), password protected and accessible only to Statespace employees conducting blinded data analyses. Data were accessed for analysis via structured query language (SQL) scripts that extract in-game data (e.g., x- and y-cursor positions, screen resolution and framerate, target locations, target presentation times and durations, etc.) and other electronically – collected data. No personally identifying information was acquired. To associate data from each round with an individual player, the software generated a unique alphanumeric identifier for each player, which was kept only to associate data analyzed for the study with the full dataset in the remote server.

Participants used the default mouse sensitivity settings, and each participant used the same mouse sensitivity settings for all testing days.

### Description of Playlist tasks

#### 1. Spidershot

Targets (orbs) are presented on the screen in a series of trials at pseudo-random intervals. Each inter-trial interval is drawn from a pre-specified time range. The participant must fire their weapon on the target 3 times to “kill” the target (Figure 2). The difficulty of the task is modulated by varying target size and/or target duration (the duration of time during which a target is presented). This task assesses motor control of ballistic movements by measuring the speed (proportion of distance to the target/time in milliseconds), precision, accuracy (proportion of distance to target), and reaction time of each mouse movement a participant makes, as well as the standard deviation of these measurements across multiple targets.

**Figure 2.**
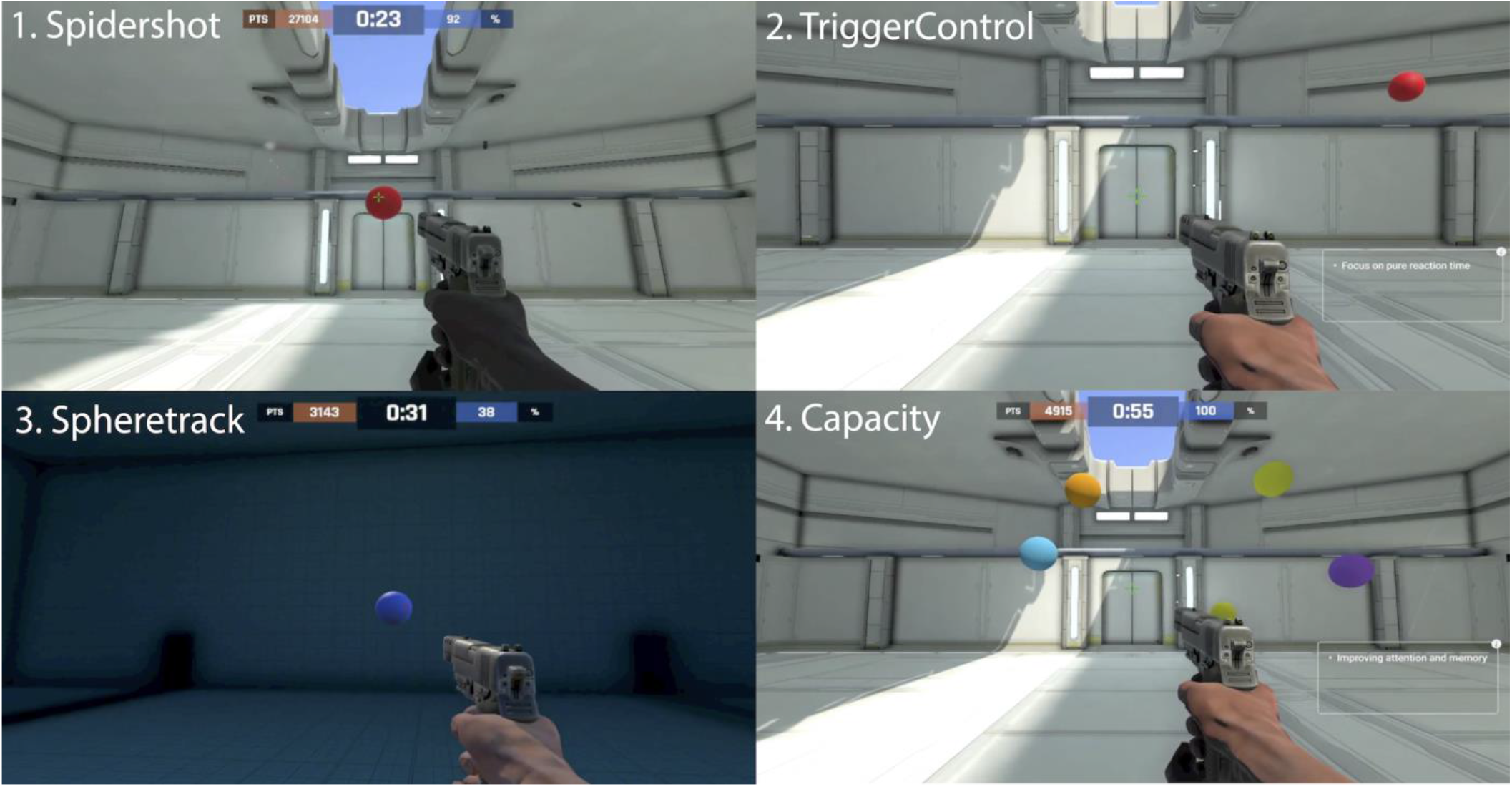
Screenshots of Playlist tasks.

#### 2. TriggerControl

This task is an embodiment of a go/no-go test implemented as a FPS shooting scenario. For each of a series of trials, a single target appears on the screen in one of two colors: blue or red. The participant is instructed to click the mouse as quickly as possible when a red target appears (Figure 2). If the target is blue, the participant is instructed to withhold the mouse click. A sound is presented synchronously with target appearance, regardless of color. Targets are presented at pseudo-random intervals. Each inter-trial interval is drawn randomly from a pre-specified time range. Difficulty of the task is modulated by the rate (the mean interval) at which new targets appear. This task assesses visual sensory processing speed and accuracy as well as cognitive control to inhibit a prepotent response, by proportion of hits (mouse clicks to the correct color), false alarms (mouse clicks to the incorrect color), and reaction time for hits and false alarms.

#### 3. Spheretrack

With their virtual weapon, the participant must track or follow a target as it moves at random in the virtual environment, keeping the crosshair on the target orb for as long as possible (Figure 2). Difficulty of the task is modulated by the speed and predictability of target motion. This task assesses motor control of tracking movements and visual sensory processing of motion by measuring the proportion of time that the participant is on versus off target.

#### 4. Capacity

For each of a series of trials, multiple targets appear on the screen at once in different colors (Figure 2). The targets disappear briefly and when they reappear, one has changed color. The participant must identify and shoot the target that has changed color. Colors are chosen from a colorblind-friendly palette. Difficulty of the task is modulated by the number of targets that appear on screen, increasing by 1 after a correct response and decreasing by 1 after an incorrect response. This task assesses visuo-spatial working memory by measuring the proportion of trials in which the participant hits the correct target that has changed color and the maximum number of targets on screen for which the participant hits the correct target. Specifically, a participant’s performance is used to calculate a “Visual Capacity Threshold”, which corresponds to the number of orbs that lead to 50% correct responses on average.

### Gaming Hardware

All hardware for the gamified assessment is commercially available and non-invasive: AsusTUF Gaming Laptop 15.6” Full HD IPS, 8th-Gen Intel Core i5-8300H, GTX 1050 4GB, 8GB DDR4, 256GB M.2 SSD, Gigabit WiFi - FX504GD-WH51 (AsusTek Computer Inc, Taipei, Taiwan), VicTsing wired mouse model SC170828US (VicTsing Technology Co., Ltd, Sunnyvale, CA), Dodocool Stereo Gaming Headset B079P2VQBG for PS4 PC, 7.1 Surround Sound Over Ear Headphones with Noise Cancelling Built-in Mic, LED Light, Soft Memory Earmuffs for Laptop Max, USB Interface (Dodocool, HongKong).

### Data/Statistical Analyses

All Visual Analog Scale measures, heart rate, and blood pressure were entered into two separate Microsoft Excel spreadsheets (i.e., manual double-key data entry) and compared to assure data quality prior to analysis. Aim Lab playlist data were automatically recorded through Aim Lab software and stored on a secure Amazon Web Services account. Blinded statistical analyses were performed in the R statistical programming environment.

All analyses, other than ANCOVA, were performed on data after normalizing by first subtracting the baseline score/marker at time 0 (pre-treatment) from the subsequent scores acquired at 60 min, 120 min, and 180 min post treatment. Absolute change from baseline has higher power than percentage change from baseline^27^.

Assumptions for ANOVA were tested using functions in the rstatix package in R^28^. For each measure, outliers were checked via the boxplot method. Values above Q3 + 1.5xIQR or below Q1 - 1.5xIQR were considered outliers and values above Q3 + 3xIQR or below Q1 - 3xIQR were considered extreme outliers. Normality assumptions were checked on all variables using a one-sample Shapiro-Wilk test. Levene’s test was used to assess the assumption of homogeneity of variance across groups.

Because assumptions of normality and outliers were not met, a two-way between-within subjects (PL vs. CAFF vs. CMT X 60 min, 120 min, 180 min) repeated measures ANOVA on change from baseline at 0 min was performed using trimmed means with the WRS2 package, bwtrim function in R. This method is robust to deviations from assumptions required for traditional ANOVA without transforming data and using trimmed means obviates the need to remove outliers^29^.

Post-hoc tests for differences between treatment group medians were performed using the mcp2a function in the WRS2 package in R, corrected for multiple testing within each measure using the family-wise error rate (FWE), using percentile bootstrap for confidence intervals and p-values. To minimize multiple testing issues, post-hoc tests were only performed for measures showing significant main effects (p < 0.05) or for those suggestive of significance (0.05 ≤ p < 0.10) from the ANOVA.

Effect sizes were calculated using the t1way function in the WRS2 package in R. This method uses an explanatory measure of effect size ξ^30^. Values of ξ = 0.10, 0.30, and 0.50 correspond to small, medium, and large effect sizes. It is a robust generalization of Cohen’s d and can be applied to more than two groups/trials.

ANCOVA was performed, correcting for scores at time 0, to evaluate if significant effects were dependent on differences in starting scores between treatments. ANCOVA analysis was performed in R for Aim Lab playlist markers that were found to be statistically significant in post-hoc testing.

When multiple tests are not independent, which is the case for many of the variables within and between Aim Lab tasks as well as among the three between-group comparisons in post hoc tests; it is appropriate to use a less stringent correction method for multiple testing^31^. Thus, FWE was used to correct for multiple testing when required.

## Results

### Aim Lab playlist markers

Assumptions of normality and no outliers were violated for multiple outcome variables, thus techniques used were robust to violations of assumptions for ANOVA. Assumption of homogeneity across groups was not violated for any variable.

All results reported below were normalized by first subtracting the baseline score/marker (pre-ingestion of the supplement) from the subsequent scores that were acquired during each visit. Consequently, when we report a statistically significant change in performance for one treatment over another, it should be interpreted as a significantly larger change in performance from baseline.

### ANOVA

Using a robust two-way repeated measures ANOVA on change from baseline, no significant treatment X time interaction effects were found for any Aim Lab task playlist marker. No significant main effects were found for session time for any Aim Lab task playlist marker. Four Aim Lab task playlist markers showed statistically significant main effects for treatment (Trigger control: median reaction time false alarms (p = 0.002); Spheretrack: proportion time on (p = 0.005); Spidershot: median kill time (p = 0.008); and Capacity: visual capacity threshold (p = 0.029) and three were suggestive of significance (Trigger control: median reaction time hits (p = 0.054); Spidershot: reaction time standard deviation (p = 0.055); Capacity: median time to kill hits (p = 0.087) (Figure 3).

**Figure 3.**
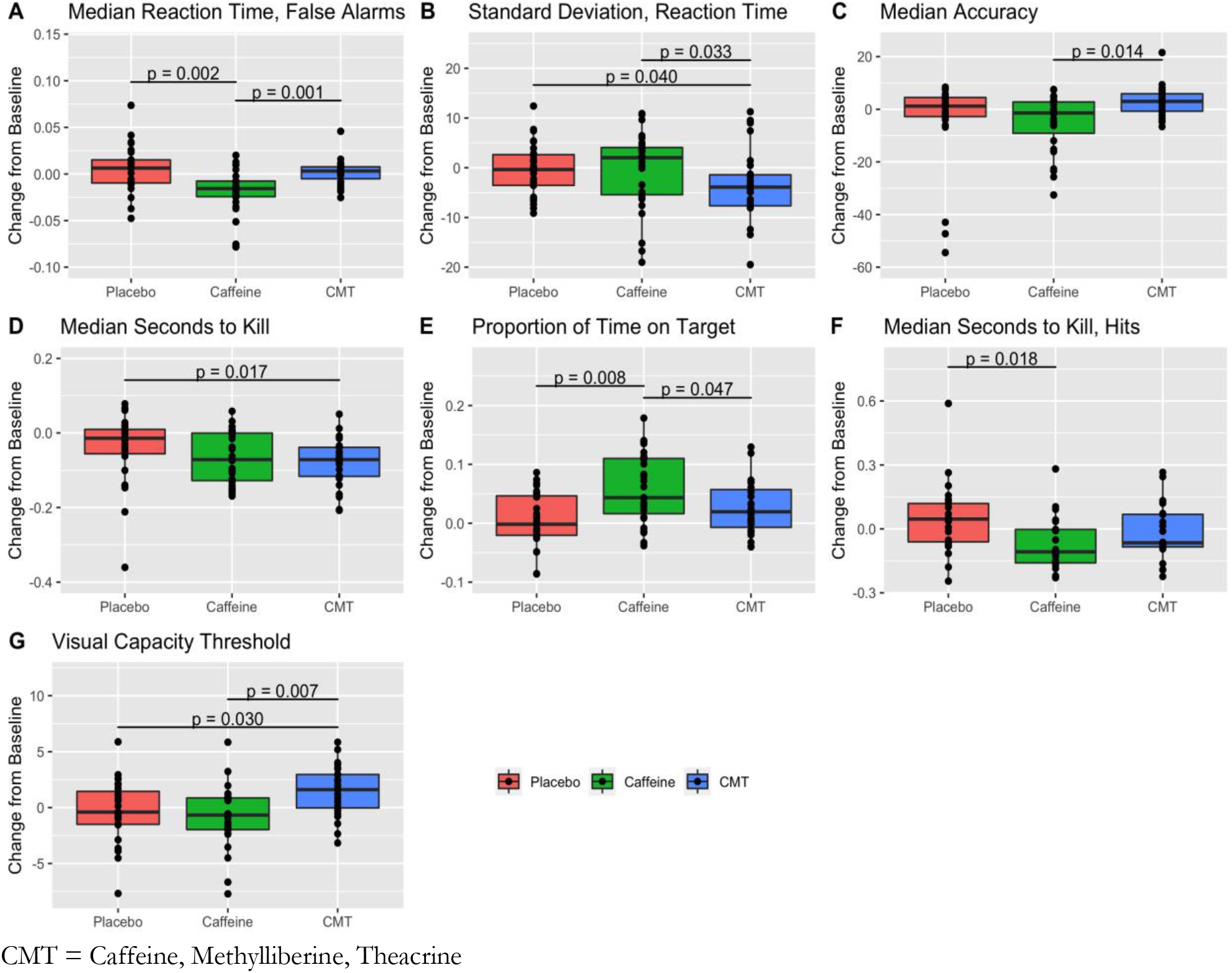
Boxplots of significant (p≤0.05) and suggestive of significant (p<0.10) Aim Lab task playlist markers.

### Post-hoc Main Effect Contrasts

Of the playlist markers that showed statistically significant and suggestive of significant main effects for treatment, post hoc main effect contrasts showed statistically significant differences between treatment type for six of the seven. One post hoc marker showed differences suggestive of significance (Table 2).

**Table 1:**
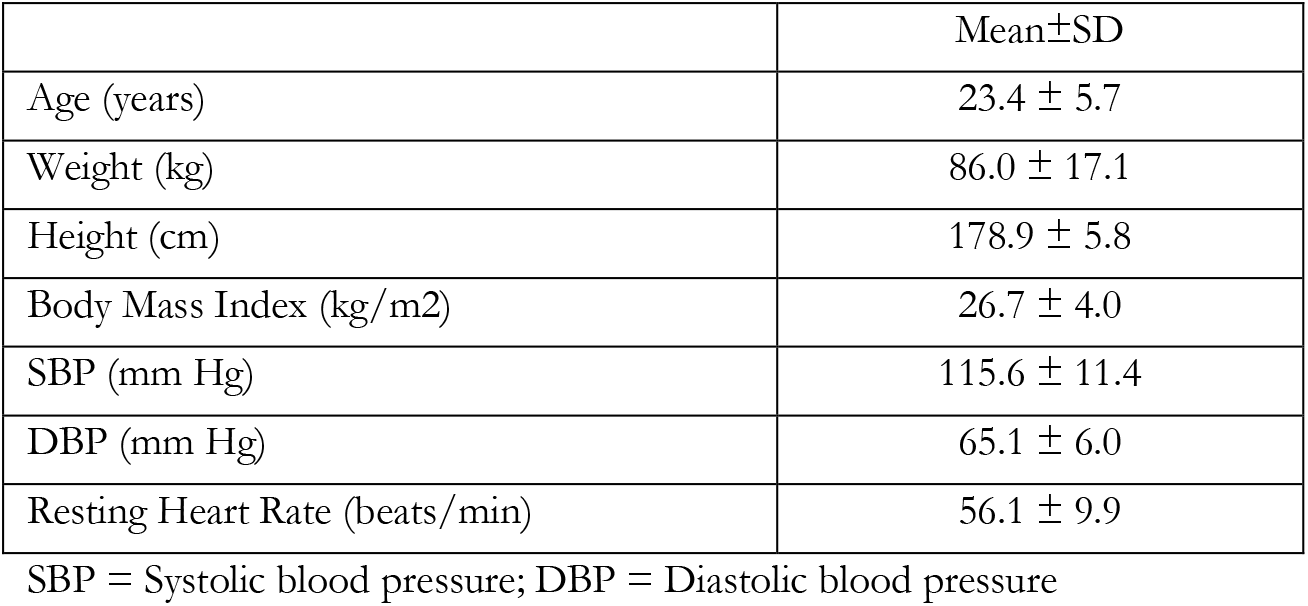
Baseline characteristics of participants (n = 9).

**Table 2:** Post-hoc between-treatment differences in median change in score from baseline for Aim Lab playlist markers that showed statistically significant and suggestive of significant main effects for treatment, p-values corrected for multiple testing using family-wise error rate (FWE), ξ effect size and the confidence interval for effect size. Effect sizes were calculated only for variables with a statistically significant between-treatment difference between median change from baseline.

| task | variable | groups | groups diff | groups diff p-value | PL median change from baseline | CAFF median change from baseline | CMT median change from baseline | $\xi$ effect size | $\xi$ p-value | 95% CI |
| --- | --- | --- | --- | --- | --- | --- | --- | --- | --- | --- |
| spidershot | median killTime | PL-CMT | 0.057 | 0.017* | -0.014 | -0.071 | -0.071 | 0.35 | 0.02 | 0.07 - 0.61 |
| spidershot | median killTime | PL-CAFF | 0.057 | 0.061 | -0.014 | -0.071 | -0.071 | NA | 0.02 | NA |
| triggercontrol | median reactionTime falseAlarms | CAFF-CMT | -0.02 | 0.001* | 0.006 | -0.017 | 0.003 | 0.61 | 0 | 0.35 - 0.87 |
| triggercontrol | median reactionTime falseAlarms | PL-CAFF | 0.023 | 0.002* | 0.006 | -0.017 | 0.003 | 0.61 | 0 | 0.35 - 0.87 |
| triggercontrol | median reactionTime hits | CAFF-CMT | -0.008 | 0.082 | -0.004 | -0.009 | -0.001 | NA | 0.13 | NA |
| capacity | median timeToKill hits | PL-CAFF | 0.164 | 0.018* | 0.036 | -0.128 | -0.075 | 0.37 | 0.04 | 0.11 - 0.61 |
| capacity | median timeToKill hits | PL-CMT | 0.111 | 0.065 | 0.036 | -0.128 | -0.075 | NA | 0.04 | NA |
| spheretrack | Proportion TimeOn | PL-CAFF | -0.046 | 0.008* | -0.002 | 0.044 | 0.019 | 0.5 | 0.01 | 0.18 - 0.75 |
| spheretrack | Proportion TimeOn | CAFF-CMT | 0.025 | 0.047* | -0.002 | 0.044 | 0.019 | 0.5 | 0.01 | 0.18 - 0.75 |
| spidershot | spar reaction stdev | CAFF-CMT | 6.291 | 0.033* | -0.357 | 2.029 | -4.262 | 0.37 | 0.01 | 0.14 - 0.64 |
| spidershot | spar reaction stdev | PL-CMT | 3.905 | 0.04* | -0.357 | 2.029 | -4.262 | 0.37 | 0.01 | 0.14 - 0.64 |
| capacity | visual capacity threshold | CAFF-CMT | -2.277 | 0.007* | -0.403 | -0.675 | 1.602 | 0.48 | 0.01 | 0.3 - 0.75 |
| capacity | visual capacity threshold | PL-CMT | -2.005 | 0.03* | -0.403 | -0.675 | 1.602 | 0.48 | 0.01 | 0.3 - 0.75 |

PL = Placebo; CAFF = Caffeine; CMT = Caffeine, Methylloberine, Theacrine. All time-based measurements (reaction time or time to kill) are measured in seconds; proportion time on is ratio; visual capacity threshold is measured in N targets. \*Significant p-value.

Compared to CAFF (p = 0.007) and compared to PL (p = 0.030), CMT was associated with a statistically significant greater improvement of participants’ Visual Capacity Threshold in the Capacity task, a measure of visuo-spatial working memory, from baseline. The effect size was medium (ξ = 0.48), as measured by Wilcox and Tian’s effect size. There was a statistically significant greater decrease in median time to kill for correctly identified targets in the Capacity task for CAFF compared to PL (p = 0.018, ξ = 0.37) and a non-significant but suggestive greater decrease (p = 0.065, ξ = 0.37) for CMT compared to PL.

Compared to PL, CMT was associated with a statistically significant greater decrease in median kill time in the Spidershot task (p = 0.017, ξ = 0.35), while CAFF had a non-significant but suggestive greater decrease in median kill time compared to PL (p = 0.061, ξ = 0.35). Compared to CAFF (p = 0.033, ξ = 0.37) and compared to PL (p = 0.04, ξ = 0.37), CMT was associated with a statistically significant greater decrease from baseline in the standard deviation of reaction time in the Spidershot task.

There was a statistically significant greater increase from baseline for the Spheretrack task proportion of time on target for CAFF compared to PL (p = 0.008, ξ = 0.50) and CAFF compared to CMT (p = 0.047, ξ = 0.50). Compared to both PL (p = 0.002, ξ = 0.61) and to CMT (p = 0.001, ξ = 0.61), CAFF, was associated with a statistically significant greater decrease in median reaction time for false alarms for the Trigger control task.

ANCOVA results show that after adjustment for measure at baseline, there was no statistically significant interaction between time and intervention type on change from baseline for the outcome measures found to have significant main effects for intervention, indicating that the effect of intervention type on score does not depend on the starting score and vice versa.

### Vitals and VAS measurements

The compounds were safe when administered together with no adverse events (Table 3).

**Table 3:** Post-hoc between-treatment differences in median change in score from baseline for all physiological and VAS variables, p-values corrected for multiple testing using family-wise error rate (FWE), ξ effect size and the confidence interval for effect size. Effect sizes were calculated only for variables with a statistically significant between-treatment difference between median change from baseline.

| variable | groups | groups diff | PL median change from baseline | CAFF median change from baseline | CMT median change from baseline | median diff p-value | $\xi$ effect size | $\xi$ p-value | 95% CI |
| --- | --- | --- | --- | --- | --- | --- | --- | --- | --- |
| SBP change | PL-CMT | -6 | -4 | 6 | 2 | 0.0202* | 0.42 | 0.03 | 0.17 - 0.61 |
| SBP change | PL-CAFF | -10 | -4 | 6 | 2 | 0.0254* | 0.42 | 0.03 | 0.17 - 0.61 |
| performance assessment change | PL-CMT | -1 | 0.8 | 0.5 | 1.8 | 0.0354* | 0.35 | 0.05 | 0.17 - 0.63 |
| jitteriness change | PL-CAFF | -1 | 0 | 1 | 0.5 | 0.0434* | 0.27 | 0.09 | 0.1 - 0.58 |
| alertness change | PL-CMT | -1 | 0.2 | 0.5 | 1.2 | 0.0874 | 0.29 | 0.27 | 0.06 - 0.59 |
| Decision making change | PL-CMT | -0.7 | -0.1 | 0.7 | 0.6 | 0.0928 | 0.23 | 0.26 | 0.08 - 0.52 |
| Decision making change | PL-CAFF | -0.8 | -0.1 | 0.7 | 0.6 | 0.094 | 0.23 | 0.26 | 0.08 - 0.52 |

PL = Placebo; CAFF = Caffeine; CMT = Caffeine, Methylliberine, Theacrine; SBP = Systolic Blood Pressure in mmHg; All visual analog scale (VAS) measures are in millimeters. \*Significant p-value.

Assumption of normality was violated for multiple outcome variables and assumption of no outliers was violated for one variable, thus techniques used were robust to violations of assumptions for ANOVA. Assumption of homogeneity across groups was not violated for any variable.

### ANOVA

Using a robust two-way repeated measures ANOVA on change from baseline, no significant treatment X time interaction effects or main effects for time were found for VAS variables (energy, alertness, focus, creativity, decision making, processing speed, irritability, jitteriness, assessment of gaming performance) or for heart rate and blood pressure. Two markers had significant main effects for treatment [SBP (p = 0.012); performance assessment (p = 0.016)] and three were suggestive of significance [jitteriness (p = 0.07); decision making (p = 0.083); alertness (p = 0.095)].

### Post-hoc Main Effect Contrasts

Of the five markers that met the threshold for significance or were suggestive of significance in ANOVA, post hoc main effect contrasts showed significant differences between treatment type for three of the markers. There was a statistically significant greater increase from baseline in SBP for CMT compared to PL (p = 0.020) and for CAFF to PL (p = 0.025): explanatory measure of effect size ξ = 0.37. Based on VAS questions, CAFF compared to PL was statistically significantly associated with a greater increase from baseline in jitteriness (p = 0.043, ξ = 0.27) while CMT compared to PL was associated with a statistically significant increase from baseline in performance self-assessment (p = 0.035, ξ = 0.35) (Table 3).

ANCOVA results showed that after adjustment for measure at baseline, there was no statistically significant interaction between time and intervention type on change from baseline for the outcome measures found to have significant main effects for intervention, indicating that the effect of intervention type on score does not depend on the starting score and vice versa.

## Discussion

The objectives of this study were to measure and compare changes from baseline in cognitive and motor performance during a FPS gaming scenario along with subjective feelings in energy, alertness, focus, creativity, decision making ability, processing speed, irritability, and jitteriness, and to assess safety and tolerability. CMT appeared to have a greater positive change from baseline in working memory over CAFF and PL and a greater positive change from baseline in speed compared with PL. In addition, CMT had a greater positive impact on cognitive control and consistency of players’ reaction time, suggestive of augmented player performance. On the other hand, CAFF appeared to have a greater positive change in motor control compared with CMT and PL. Further, CAFF led to a greater positive impact on tasks where speed was needed, but at the same time had a greater negative change in participants’ reaction time on false positives during Trigger control, indicative of a decrement in player performance. CMT and CAFF had a greater change from baseline on increasing systolic blood pressure compared with PL while remaining within normal clinical limits. Lastly, CMT had a greater impact on the change in the player’s self-assessed performance after each gaming session versus PL, while CAFF suggested a greater increase in jitteriness over PL. From a clinical perspective, the compounds were safe when administered together with no adverse events.

The results demonstrate that CMT improved cognitive abilities and speed from baseline as measured by visuo-spatial working memory in the Capacity task (i.e., visual capacity threshold) as well as median kill time in the Spidershot task. Alternatively, CAFF improved reaction time from baseline to the appearance of a target on screen and speed of mouse movement, but this was accompanied by decreased player reliability or precision in the form of a larger standard deviation from baseline in comparison to CMT. A larger standard deviation from baseline may negatively impact esports gamer performance when a high degree of precision is required under high-pressure conditions in first-player shooter and marksmanship games^32^. In addition, CAFF decreased cognitive control as measured by decreased reaction time to false alarm targets from baseline (i.e., the player became trigger-happy) in the Trigger control task and increased self-reported feelings of jitteriness. These findings parallel Rodgers et al.^11^ that caffeine made them faster not smarter and caffeine’s tendency to elevate feelings of jitteriness. Results also suggest that Dynamine™ and TeaCrine® may add a potential benefit when added to a caffeinated supplement due to a dampening effect on caffeine’s positive effects on speed and reaction time while combating the negative effects of caffeine (decreased cognitive abilities & performance reliability, increased jitteriness).

Caffeine has been shown to increase focus whether it was administered alone or paired with Dynamine™ and/or TeaCrine®^23^. Additionally, caffeine and Dynamine™ have been shown to have a positive effect on being attentive^23^ which may positively influence visuo-spatial working memory. Even though the current investigation did not find any differences in energy, alertness, or focus it did find that participants rated their overall gaming performance higher over the course of their CMT trial compared with PL. Therefore, the participants may not have felt more energetic, alert, or focused, but they felt as though they performed better overall. It seems possible that TeaCrine® may have offered a calming effect as CMT did not show a significant elevation in jitteriness contrary to CAFF^18,21^. Nevertheless, caffeine with Dynamine™ and TeaCrine® may offer a benefit to caffeine sensitive consumers to obtain cognitive enhancements without the negative effects of CAFF.

Although SBP transiently, but significantly, increased in both caffeine containing supplements, there does not appear to be any long-term effects on blood pressure^24^. The significant increase from baseline in SBP associated with either caffeine containing supplement compared to PL was consistent with Bloomer et al.^23^ showing that Dynamine™ and TeaCrine® do not have any influence on caffeine’s acute effect on SBP. However, in contrast to Bloomer et al.^23^, the caffeine containing supplements in the current investigation did not have a significant impact on DBP. Therefore, despite the acute rise in SBP subjects felt as if they were performing better with CMT rather than having an increased feeling of jitteriness with CAFF.

We acknowledge our study used a relatively small homogenous sample size (i.e., 9 young men). Future studies should include a larger, more diverse sample and should investigate long-term supplementation and cognitive/gaming performance over time (i.e., 4 weeks of supplementation with gaming sessions at pre, mid, post). In addition, the effects of various dosages of caffeine combined with Dynamine™ and TeaCrine® would help identify if an optimal mixture exists for cognitive benefits.

This investigation found CMT confers an advantage in enhancing psychomotor performance over CAFF and PL in a FPS gaming scenario. The benefits seen with CMT during gaming, as a change from baseline, were increases in working memory, speed, cognitive control, greater consistency in reaction time, and a positive self-assessment which falls in line with preferred gamer attributes^4^,^5^. Alternatively, CAFF did provide a slight advantage over CMT and PL in tracking an object and potentially identifying a target on screen by moving the mouse more quickly; however, the increased jitteriness led to an overreaction to false targets. Even though CMT and CAFF acutely increased SBP, measures were well within normal clinical limits and both were well tolerated with no adverse events reported. These data support the addition of TeaCrine® and Dynamine™ to caffeine in gamers looking to improve overall psychomotor performance during FPS gaming.

## Media-Friendly Summary

This study used AI-driven Aim Lab software developed by a team of neuroscientists to objectively assess and compare speed, precision, accuracy, reaction time, working memory capacity, and cognitive control during a first person-shooter (FPS) gaming scenario. The results demonstrated that supplementation with 75 mg Dynamine™ and 50 mg TeaCrine® amplified key aspects of cognitive performance such as working memory, speed, cognitive control, and greater consistency in reaction time when added to a moderate 125 mg dose of caffeine (i.e., the amount found in an 8 oz cup of coffee). In addition, Dynamine™ plus TeaCrine® were able to decrease the error rates (e.g., identifying false targets in FPS scenario) and jitteriness associated with isolated caffeine supplementation. Self-assessment of performance was also highest in the group using Dynamine™ plus TeaCrine® and caffeine. These data support the addition of TeaCrine® and Dynamine™ to caffeine in recreational gamers looking to improve their performance in esports.

## Author Contributions

DH assisted with study design and manuscript preparation. ID carried out data analyses and assisted with study design and manuscript preparation. JL carried out data analysis and manuscript preparation. WM assisted with study design. TJ assisted with study design and coordination. ML carried out participant recruitment, data collection, coordination of the study and compliance., assisted with data analysis, interpretation, and prepared the manuscript. All authors read and approved the final manuscript.

## Acknowledgments

The authors are grateful and recognize the contributions made by Statespace and Aim Lab towards data collection on this project, and Hector L. Lopez, M.D. and Tim N. Ziegenfuss, Ph.D. for their contributions to the study design. The authors would like to thank all the study participants who completed the study protocol. Publication of these results should not be considered as an endorsement of any product used in this study by the Center for Applied Health Sciences, Statespace, Aim Lab, or any of the organizations where the authors are affiliated.

## Availability of Data and Materials

The data and materials for this manuscript are not scheduled to be made publicly available due to the proprietary nature of the investigated materials. Contractually, the data are owned by Compound Solutions, Inc.

